# Nanopore-based native RNA sequencing of human transcriptomes reveals the complexity of mRNA modifications and crosstalk between RNA regulatory features

**DOI:** 10.1101/2024.07.11.603105

**Authors:** Yerin Kim, Kieran O’Neill, Jean-Michel Garant, Luke Saville, Simon Haile, Maryam Ghashghaei, Yongjin P. Park, Steven J.M. Jones, Ly P Vu

## Abstract

The identification and functional characterization of chemical modifications on an mRNA molecule, in particular N^6^-methyladenosine (m^6^A) modification, significantly broadened our understanding of RNA function and regulation. While interactions between RNA modifications and other RNA features have been proposed, direct evidence showing correlation is limited. Here, using Oxford Nanopore long-read direct RNA sequencing (dRNA-seq), we simultaneously interrogate the transcriptome and epitranscriptome of a human leukemia cell line to investigate the correlation between m^6^A modifications, mRNA abundance, polyA tail length and alternative splicing. We demonstrated that high quality dRNA-seq is critically important for unbiased and large-scale correlative analyses of these RNA features. The genome wide landscape of RNA methylation was captured with single nucleotide and individual isoform resolution. The length of polyA tails were ascribed to individual transcripts and genes, allowing for unprecedent measurement of polyA tail length distribution. Global analysis indicated negative association between polyA tail length and mRNA abundance while uncovering pathway-specific responses upon depletion of m^6^A forming enzyme METTL3. Overall, our study presented a rich dRNA-seq data resource which has been validated and can be further exploited to inquire into the complexity of RNA modifications and potential interplays between RNA regulatory elements.

## Introduction

RNA is a versatile molecule, which can be diversely and dynamically processed and modified. The emerging field of epitranscriptomics revealed a fundamental layer of regulation of the transcriptome with identification and characterization of a number of chemical modifications decorating the nucleobases of a messenger RNA (mRNA)^1^. The classical and most well studied modifications on mRNAs are the N7-methylguanosine (m7G) 5’ cap and polyadenylation at the 3’ end^2^. The most abundant internal modification in eukaryotic mRNAs is the N^6^-methyladenosine (m^6^A) modification. m^6^A modifications are predominantly deposited by the methyltransferase “writer” complex composed of the catalytic heterodimer METTL3/METTL14 and can be removed by two major “eraser” enzymes: FTO and ALKBH5. m^6^A sites are recognized and interpreted by “reader” proteins including YT521-B homology (YTH) family of reader proteins (YTHDC1/2 and YTHDF1/2/3)^3^. RNA methylation plays a key role in modulating many critical physiological processes such as control of stem cell function and differentiation^4, 5^, immunity^6, 7^ and cancer development^8, 9^. These highlight the importance of precise identification and characterization of RNA modifications in the context of their interaction with various RNA features to influence the fate of an mRNA during its life cycle.

Development of next generation sequencing (NGS) – based methods including m^6^A-specific methylated RNA immunoprecipitation with next-generation sequencing (MeRIP-Seq)^10^, m^6^A individual-nucleotide-resolution cross-linking and immunoprecipitation (miCLIP-seq) and evolved TadA-assisted N^6^-methyladenosine sequencing (eTAM-seq)^11^ have enabled profiling of m^6^A distributions across transcriptomes in an unbiased and genome wide manner. Examination of the biological consequences of m^6^A on target genes can be performed by incorporating m^6^A profiles with assessments of other features of the transcriptome. For example, integrating of me-RIP seq data and Photoactivatable-Ribonucleoside-Enhanced Cross linking and Immunoprecipitation (PAR-CLIP) to map YTHDF2’s bound mRNAs identified m^6^A modified transcripts susceptible to mRNA decay mediated by YTHDF2^12^. YTHDF2 can recruit the CCR4-NOT complex, which in turn deadenylates mRNAs and initiates mRNA degradation^13^. Combined m^6^A profiling and Ribosome profiling revealed m^6^A dependent translation control over a subset of target genes^8, 14^. To date, it is known that the presence of m^6^A modifications on an mRNA can influence how it is spliced^15^, exported^16^, degraded^12^ and translated^14, 17^. Notably, transcriptional activity^18^ and the interactions between mRNAs and spliceosomes can affect deposition of RNA modifications^19, 20^, indicating dynamic interactions between different RNA processing mechanisms.

While these studies provided foundational knowledge of the function and biological significance of RNA modifications, the associations between m^6^A methylation and other RNA features were inferred using independent techniques and datasets obtained from different experimental settings or biological samples. Therefore, genome wide correlation of these elements is still inadequate or completely missing. Even though a potential interaction between m6A modifications and polyA tail length can be speculated from recruitment of the CCR4-NOT complex to the modified transcripts by YTHDF2^13^, the actual evidence of interaction is not available. This is mainly due to challenges measuring polyA tail length using specialized methods^21^ and to challenges with capturing it simultaneously with m^6^A profiling. In addition, the lack of stoichiometric measurement of modified vs. unmodified RNAs limits comparison of RNA features including mRNA expression level, mRNA decay and translation activity between non-modified transcripts and modified transcripts. Thus, it is not well established whether the different rate of RNA modifications can result in different level of changes in these parameters. Expanding the characterization landscape of RNA modifications and reducing their overall bias will enable a more holistic understanding of how RNA modifications influence mRNA function and regulation.

Oxford Nanopore long read direct RNA sequencing (dRNA-seq) has recently emerged as an alternative approach to short-read NGS methods enabling the characterization of mRNAs by sequencing the native RNA molecule, and thereby preserving the chemical modifications decorating their nucleobases for sequencing^22–25^. dRNA-seq identifies bases i.e. adenine (A), uracil (T/U), cytosine (C), and guanine (G) and their respective base modifications based on the measurement of current intensity signatures corresponding to each base and current shifts when an RNA molecule passes through protein pores. Moreover, the long-read capacity of dRNA-seq provides measurement of RNA abundance at both the gene and the transcript (isoform) levels. Without the use of several specialized methods, dRNA-seq simultaneously measures many RNA modifications including m^6^A RNA methylation,^23^ and the polyA tail^26^. Importantly, detection of the ratio of altered signals vs. standard signals provides quantitative measurement of RNA modification frequencies at single nucleotide resolution in individual transcripts^27^. dRNA-seq has the potential to bring unprecedented insights into the complexity of the transcriptome and epitranscriptome. However, requirements for high quality datasets have been a barrier to optimally exploit the technology. The majority of available dRNA-seq datasets in human cell lines were obtained using the MinION platform, whose outputs are about 1 million reads/sample on one flow cell run, detecting less than a thousand unique transcripts and representing a small proportion of a full human transcriptome^23, 28^. The inadequate coverage, thus, limits genome wide correlation analyses between different RNA modifications and features. Here, we performed dRNA-seq from the MOLM13 leukemia cell line using the high throughput PromethION platform to produce rich datasets with the depth and coverage necessary to assess gene expression and RNA modification dynamics. MOLM13 is a well-established cell line model for the study of leukemia and it has previously been assessed for its genome wide m^6^A distribution using orthogonal techniques^8, 29^. Here, we gained additional insights into the interplay of m^6^A posttranscriptional modifications on mRNAs with other posttranscriptional modifications, such as 3’ polyadenylation and alternative splicing/ isoform usage with potential implications for leukemia biology. We also examine how depletion of m^6^A writer METTL3 in MOLM13 cells and consequently how acute reduction of m^6^A modifications, influences the biochemistry and expression levels of mRNA transcripts. Using high-quality dRNA-seq datasets, we also explored critical features of mRNA including mRNA abundance, polyA tail length and alternative splicing.

## Results

### Comprehensive capture of the transcriptome and epitranscriptome of leukemia cells using Oxford Nanopore long-read direct RNA sequencing (dRNA-seq)

We isolated poly(A) enriched mRNAs from leukemia MOLM13 cells and subjected them to sequencing using the PromethION platform for its high throughput. To explore the interaction between m^6^A RNA methylation and other RNA features, we performed the experiments using four biological repeats of control (transduced with scramble shRNA) and two biological replicates of METTL3-shRNA-mediated knocked down cells. Data generated from the platform was then processed through a standard base calling pipeline and alignment was performed using Minimap2^30, 31^ and signal segmentation was performed via f5c ‘eventalign’^32^. Further analysis was conducted to obtain information on poly(A) tail length; isoforms and m^6^A RNA methylation (**Figure 1A**). For 6 samples, we obtained over 34 million reads in total with a range from 3 to 7 million reads per sample and basecalling Phred were scored mainly above 8 **(Figure S1A-B,S1K-L)**. A substantial portion of transcripts were captured with a coverage higher than 10x, while about 30% mapped to transcripts with a coverage higher than 30x. The majority of reads are longer than 800 bp and exhibit basecalling Phred scores higher than 10 **(Figure S1C-J, S1M-T)**. These results demonstrate the datasets are of high quality and provide better outputs and coverage compared to datasets generated by MinION or GridION dRNA-seq platforms.

**Figure 1.**
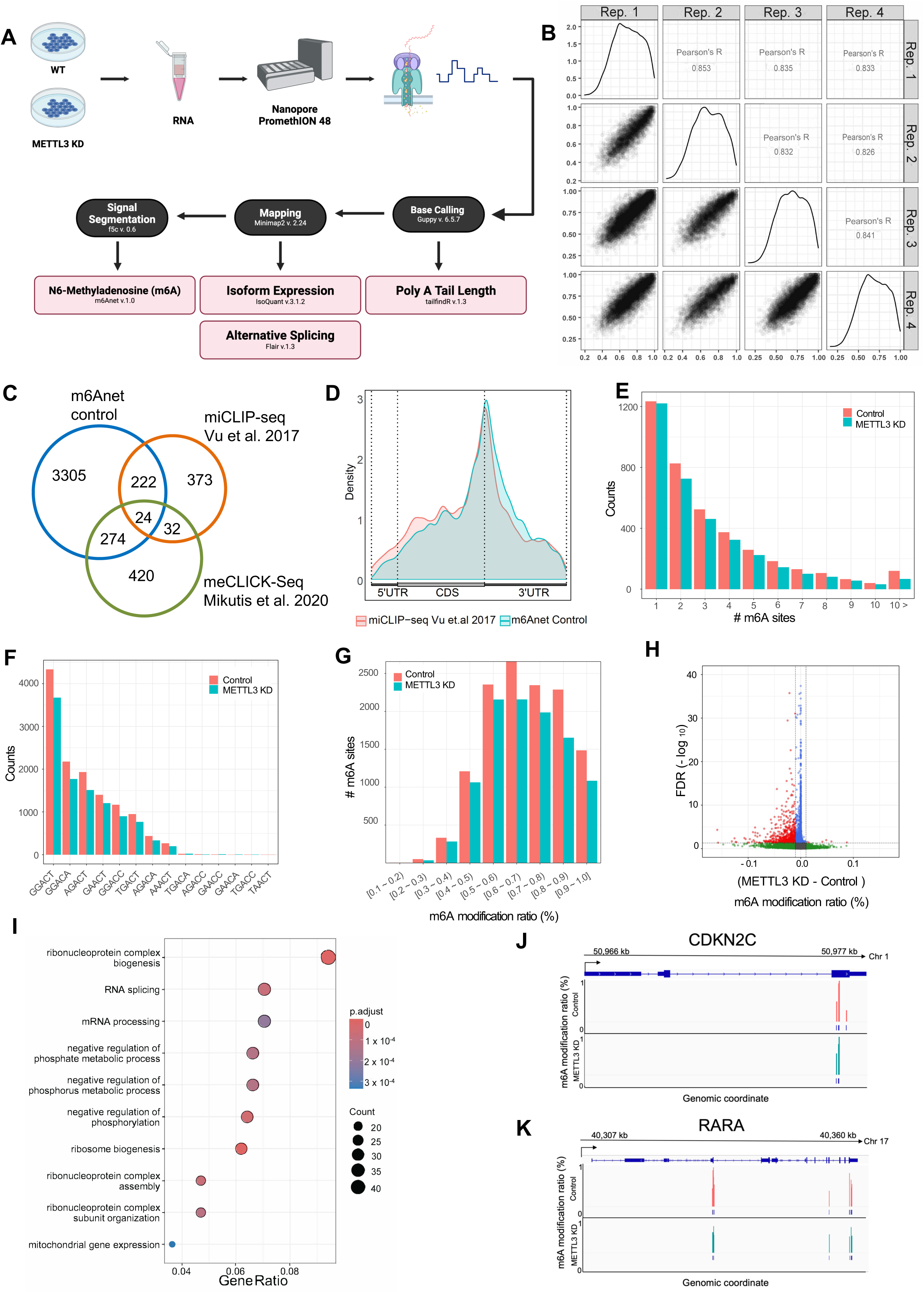
Comprehensive capture of the transcriptome and epitranscriptome of leukemia cells using Oxford Nanopore long-read direct RNA sequencing (dRNA-seq) **(A)** Workflow of Oxford Nanopore direct RNA sequencing (dRNA-seq) and *in silico* data analysis. **(B)** Pairwise scatterplots of Pearson’s correlation of m^6^A ratios for modification sites (m6Anet probability ≥ 0.9) identified in all four control replicates. **(C)** Venn diagram showing overlapping m^6^A genes identified by dRNA-seq and those identified previously using miCLIP-seq. **(D)** Metagene plot illustrating the transcriptome-wide distribution of m^6^A across 5’-untranslated regions (UTRs), coding sequences (CDS), and 3’-UTRs from dRNA-seq (m6Anet probability ≥ 0.9) and miCLIP-seq (Vu et al. 2017) respectively. **(E)** Bar graphs showing numbers of genes categorized based on number of m^6^A sites identified within the gene by dRNA-seq (m6Anet probability ≥ 0.9). **(F)** Bar graphs showing numbers of m^6^A sites characterized with the particular DRACH motifs. **(G)** Bar graphs showing numbers of m^6^A sites with corresponding m^6^A ratio (m6Anet probability ≥ 0.9). **(H)** Volcano plot of Mann-Whitney test result comparing m^6^A modification probability of each m^6^A modification site identified in MOLM13 cells transduced with control shRNA (control) vs. cells transduced with shRNA targeting METTL3 (METTL3-KD). X-axis shows median difference of m^6^A probability (|KD - Control| > 0.01, FDR < 0.05). Housekeeping genes RPS, RPL, MT genes were not included in the plot to avoid skewing of high FDR values due to their high expression levels. **(I)** Gene Ontology (GO) and pathway analysis of genes whose m^6^A frequencies were decreased upon METTL3 depletion (Mann-Whitney test, (KD - Control) < -0.05, FDR < 0.05). p-values were adjusted by Benjamini-Hochberg method with threshold p < 0.01 & q < 0.05. Top 10 pathways are shown. **(J-K)** Representative IGV view of m^6^A sites (m6Anet probability ≥ 0.7) on CDKN2C and RARA transcripts.

Because dRNA-seq enables the detection of RNA modification signals in native RNA strands, we assessed m^6^A RNA methylation within the transcriptome of MOLM13 leukemia cells using the neural-network-based m^6^A calling software m6Anet^33^. m6Anet was used because of its capability of providing single-molecule level stoichiometry of m^6^A at the site level and de novo identification of m^6^A sites without the requirement of a reference sample. Our analysis of the control samples showed high reproducibility (Pearson’s R = [0.83, 0.85], p-value < 0.05) across all 4 replicates **(Figure 1B)**. Each of the four replicates underwent separate processing, assessing m^6^A stoichiometry in reads exhibiting coverage of greater than 10 reads to allow for inclusion of reads that may exhibit varied expression between a higher number of replicates but still exhibit high probability scores for m^6^A modification. We identified 11,266; 4,748; 11,110 and 8,327 m^6^A sites from the 4 replicates respectively. Among the identified hits, 2,742 (17.1%), 350 (2.2%), 2,234 (13.9%), and 994 (6.2%) were unique to individual replicates **(Figure S2A).** We collectively pooled the m^6^A sites scored in at least 2 replicates with a high confidence threshold (m^6^Anet probability ≥ 0.9) to identify 12,716 m^6^A sites within 3825 genes **(Supplemental table 1)**. To evaluate the dRNA-seq based method against established assays, we intersected the list of m^6^A modified transcripts with previously published profiling of m^6^A modifications in MOLM13 using miCLIP-seq^8, 29^ and meCLICK-Seq^29^ and meRIP-seq^18, 34^. We found that 246 out of total 651 genes (∼37%) (Vu et al., 2017) and 298 out of total 750 genes (∼39%) (Mikutis et al., 2020) overlapped with dRNA-seq data (**Figure 1C**). Nanopore direct RNA-seq (m6Anet probability ≥ 0.9) detected 3305 additional m^6^A marked genes. At the same time, the shared targets between the two datasets are only 56 out of 651 and 750 genes, which is significantly lower than targets shared with dRNA-seq **(Figure 1C**). While more overlapped genes were seen with meRIP-seq, the percentages was fairly similar to those observed with miCLIP-seq and meCLICK-seq (**Figure S2B**) We observed a similar pattern in the distribution of m^6^A, as previously observed using orthogonal detection techniques^8, 10, 35^ where m^6^A modifications exhibited a substantial enrichment surrounding stop codons, distributed throughout the 5′-untranslated regions (UTRs), coding sequences, and 3′-UTRs **(Figure 1D)**. We applied the same analyses to METTL3 depleted samples (**Figure S2C)** and obtained equally high-quality transcriptome wide assessments of m^6^A RNA methylation in these cells (**Figure S2D-G and Supplemental table 2**). It is notable that out of 3825 and 3397 genes with m^6^A identified in control and METTL3 depleted cells, 2955 genes (over 75% m^6^A targets) were found in both conditions and similar overlapping genes with other datasets **(Figure S2E-F)**. We observed a global reduction in the number of m^6^A methylation sites observed across the spectrum (**Figure 1E**). Importantly, as m6Anet specifically identifies DRACH motifs, we further broke down the gene counts to each motif and found the GGACT motif, which is the most enriched m^6^A motif, also exhibited the greatest reduction (**Figure S2H and Figure 1F**). Taken together, these data strongly support the robustness of dRNA-seq detection of m^6^A.

An advantage of dRNA-seq characterization of m^6^A RNA methylation in comparison to immunoprecipitation-based methods is the capacity to estimate m^6^A stoichiometry by measuring the ratio of reads which are modified, on the basis of the frequency of m^6^A modified transcripts vs. total mRNA abundance. Therefore, we derived the modification ratios and observed a significant decrease in frequency of m^6^A upon acute METTL3 knockdown (**Figure S3F**). The reduction in m6A presented preferentially in groups of transcripts with higher m^6^A frequencies (**Figure 1G**). Differential m^6^A frequency analysis further confirmed the global loss of m^6^A RNA methylation upon METTL3 knockdown and identified a subset of targets whose m^6^A frequencies are significantly lower due to acute loss of METTL3 activity (Mann-Whitney test, with |Δ m^6^A ratio| > 0.01 and FDR < 0.05 -**Supplemental table 3** and **Figure 3H**). A subset of m^6^A sites exhibited small Δ m^6^A ratio but high FDR due to their high level of expression. In total, 606 differentially modified positions at DRACH motifs across a total of 510 genes passed the testing threshold. GO term analysis (Biological Pathway, Benjamini-Hochberg p.value < 0.01) demonstrated that these m^6^A genes are associated with ribonucleoprotein complex biogenesis, RNA splicing, and mRNA processing **(Figure 1I)**. These results are consistent with previously reported pathway analysis^34^. Additionally, because the predicted m^6^A sites are being compared to those from a knockdown reference dataset, we used the acceptable threshold of m^6^Anet probability ≥ 0.7 as recommended, for subsequent analysis,^33^ allowing a broader inclusion of modification sites for evaluation. Reduction in the peak signals can be observed clearly in target transcripts as illustrated for *CDKN2C* and *RARA* (**Figure 1J-K**). Overall, we demonstrate that dRNA-seq using the Oxford Nanopore platform produced high quality datasets and enabled comprehensive characterization of the m^6^A RNA methylation profile of a representative leukemia line.

### Correlation analysis of m^6^A RNA modifications and mRNA abundance

RNA methylation has been shown to influence many steps in an mRNA life cycle, particularly RNA processing and splicing, RNA stability and translation^9^. Unlike meRIP or miCLIP which only distinguish between m^6^A and non-m^6^A marked RNA molecules, dRNA-sequencing analysis enables the stoichiometric assessment of m^6^A decorated nucleobases. To gain insights into whether m^6^A stoichiometry impacts mRNA abundance, we performed correlative analysis of expression levels (transcripts per kilobase million-TPM) of individual transcripts with their corresponding m^6^A modification ratios. We found that globally there is a negligibly weak correlation between TPM vs. averaged m^6^A modification at the steady state in control cells (Spearman’s R = -0.071, p-value = 2.419 x 10^6^) (**Figure 2A**). Interestingly, breakdown to 5 groups based on m^6^A modification ratio at the ranges of [0.0 ∼ 0.2), [0.2 ∼ 0.4), [0.4 ∼ 0.6), [0.6 ∼ 0.8), and [0.8 ∼ 1.0] showed significantly lower TPM expression of transcripts in the [0.0 ∼ 0.2] group in compared to the more frequently modified transcripts (TPM=10.06 vs. 15.43, 15.21, 14.08, and 14.15) **(Figure 2B).** The observation challenges the view that m^6^A modifications generally correlate with decreased mRNA abundance. Additionally, we observed that increasing numbers of m^6^A sites are associated with a gradual decrease in gene expression with an exception for transcripts identified with more than 10 m^6^A sites (**Figure 2C**). The median TPM of each group is as follows: 26.92, 25.19, 22.13, 22.38, 21.05, 20.35, 20.93, 20.54, 18.68, 15.21, and 25.50 for groups characterized with 1,2,3,4,5,6,7,8,9,10, and greater than 10 m^6^A sites respectively. These results suggested that correlation between mRNA abundance and m^6^A at the steady state depends on both modification rates and number of modification sites.

**Figure 2.**
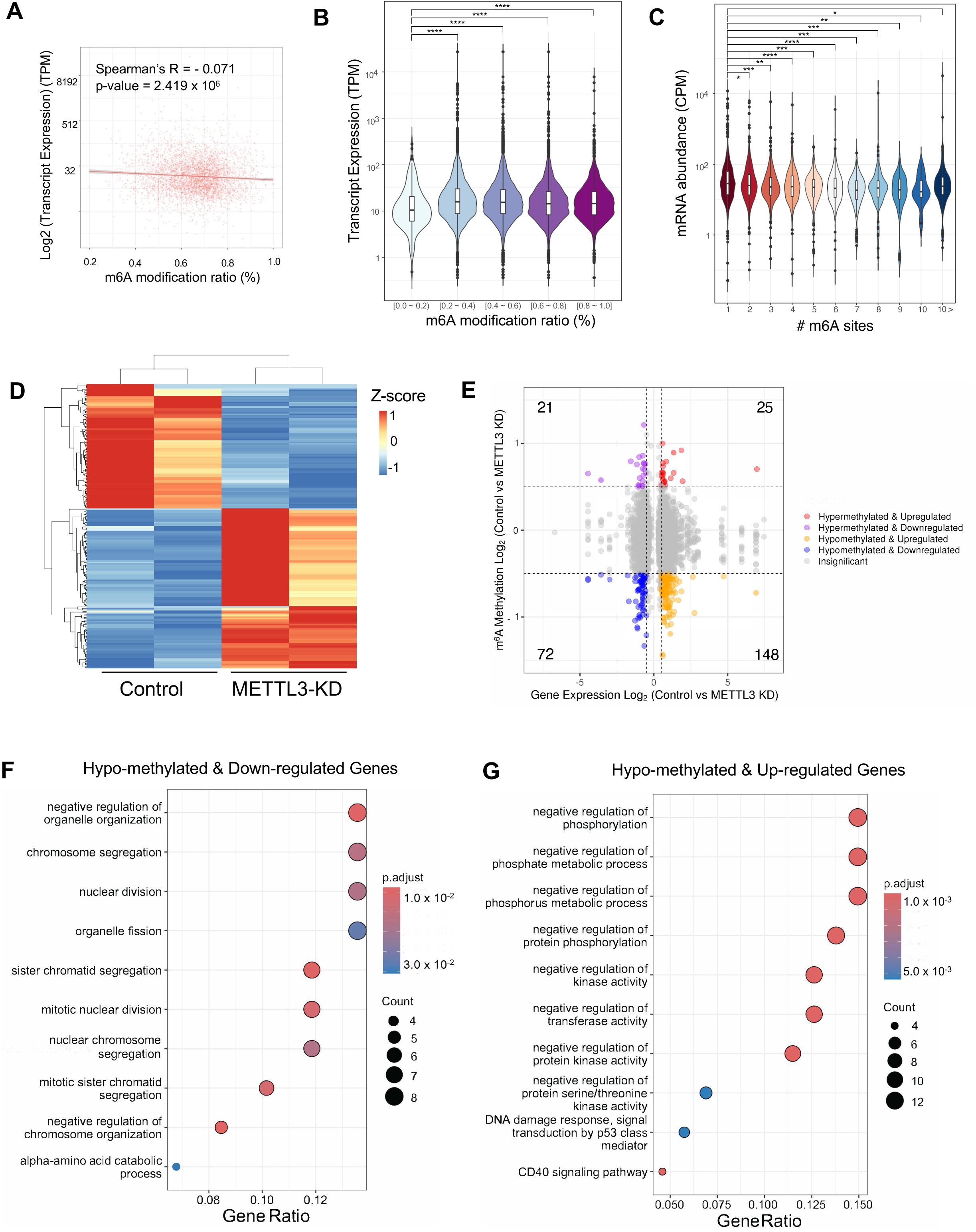
Correlation analysis of m^6^A RNA modifications and mRNA abundance. **(A)** Scatterplot showing correlation between averaged m^6^A frequencies per transcript vs. matched transcript expression (TPM) on control cells. High confident sites with m6Anet probability ≥ 0.9 sites were included in the analysis. Spearman’s R = -0.071, p-value = 2.419 x 10^-6^. **(B)** Violin plots showing transcript expression (TPM) of transcripts categorized into 5 groups of transcripts represented with increasing m^6^A modification ratio in control cells. (Wilcoxon test, * p < 0.05, ** p < 0.005, *** p < 0.0005, **** p < 0.00005) **(C)** Violin plots showing gene expression (CPM) by number of m^6^A sites (m6Anet probability ≥ 0.9) identified per gene. (Wilcoxon test, * p < 0.05, ** p < 0.005, *** p < 0.0005, **** p < 0.00005) **(D)** Heatmap showing differentially expressed genes (DEGs) in METTL3 depleted cells (METTL3 KD) vs. control. **(E)** Four quadrant diagrams showing differentially methylated sites (|Log_2_ Fold Change| ≥ 0.5, p. value < 0.05) and differentially expressed genes (|Log_2_ Fold Change| ≥ 0.5, p. value < 0.05) between control and METTL3-KD cells. Hyper-methylated & Up-regulated (hyper-up) with 20 genes and 25 m^6^A sites, Hyper-methylated & Down-regulated (hyper-down) with 20 genes and 21 m^6^A sites, Hypo-methylated & Up-regulated (hypo-up) with 97 genes and 148 m^6^A sites, and Hypo-methylated & Down-regulated (hypo-down) with 60 genes and 72 m^6^A sites. **(F-G)** GO analysis for gene groups: Hypo-methylated & Down-regulated **(F)** and Hypo-methylated & Up-regulated (**G**) p-values were adjusted by Benjamini-Hochberg method with threshold p. value < 0.01 & q. value < 0.05.

Next, we looked into the possible connection between altered m^6^A modifications and mRNA expression upon m^6^A reduction by METTL3 knockdown (METTL3 KD vs. control). Similar to previously described^8^, METTL3 depletion resulted in both up (108 genes, Log_2_ Fold Change > 1, adjusted p. value < 0.05) and down (84 genes, Log_2_ Fold Change < -1, adjusted p. value < 0.05)) regulated genes (**Figure 2D and Figure S3A and Supplemental table 4**). In agreement with reported phenotypes, we also observed that METTL3 knockdown cells exhibited increased expression of apoptotic and myeloid differentiation genes (**Figure S3B-C**). Given the simultaneous capture of differential gene expression and m^6^A modification ratios in the same samples, we performed conjoint analysis of changes in m^6^A and gene expression in METTL3 KD vs. control cells. All genes were categorized into four groups (|Log_2_ Fold Change| ≥ 0.5, p-value < 0.05), 25 hypermethylated and upregulated sites (hyper-up), 72 hypomethylated and downregulated sites (hypo-down), 21 hypermethylated and downregulated sites (hyper-down), and 148 hypomethylated and upregulated sites (hypo-up) were found **(Figure 2E, Figure S3D-G)**. It was noted that 77.42% (72/93) of downregulated mRNA transcripts and 85.55% (148/173) of upregulated mRNA transcripts were associated with m^6^A hypomethylation in KD cells. In addition, the numbers of “hypo-down” and “hypo-up” genes were more than those of “hyper-down” and “hyper-up” genes. Gene Ontology term analysis revealed that the majority of “hypo-down” genes involve in organelle organization and chromosome segregation **(Figure 2F)**. On the other hand, “hypo-up” genes associate with negative regulation of protein phosphorylation and metabolic processes **(Figure 2G)**. Taken together, these data indicated that while reduced m^6^A modifications tend to correlate with increased mRNA levels, the effect is not global and rather pathways specific.

### Genome wide characterization of poly(A) tail length and its correlation with mRNA abundance in leukemia cells

A transcript is processed and supplied with a polyadenylation (polyA) tail to protect it from RNA nuclease activity and degradation^36^. It is also known that polyA can modulate translation activity. Despite its important role in gene expression control, characterization of polyA tail length in a genome wide manner is challenging due to the repetitive nature of the tail, making it difficult to analyze in traditional short read sequencing^36^. Special assays developed to measure polyA tail lengths such as mTAIL^21^ and Poly(A)-tail profiling (PAL-seq)^37^ are arduous and often biased toward longer tailed transcripts^38^. On the other hand, dRNA-seq captures the unique streak of signals corresponding to the homopolymeric regions within the polyA tail. This enables genome wide assessment of mRNA polyA tail length at single-molecule resolution. We used Nanopolish^39^ and tailfindR^26^ to determine the lengths of polyA tails. The estimates of poly A tail length showed high reproducibility across independent control replicates (Pearson’s R = [0.70, 0.78], p-value < 0.05) **(Figure 3A and Figure S4A)**. It’s noteworthy that per-transcript estimates of polyA tail length obtained through tailfindR demonstrated a strong correlation with those acquired using Nanopolish (Spearman’s R = [0.65, 0.67], p-value < 0.05) **(Extended Data Figure 4B-C)**. The analysis provides a clear view of the full spectrum of polyA tail length distribution, ranging from a few nucleotides to close to 400 nucleotides and estimated the median length of 92.1 nucleotides for polyA tails in leukemia cells **(Figure 3B)**. Given the comprehensive and genome wide characterization of polyA tail lengths, we further categorized transcripts into three groups based on the first and third quartiles of the median polyA tail length distribution, denoted as “short,” “medium,” and “long”. The first quartile is 98, the second quartile is 116, and the third quartile is 139. Transcripts with “short” lengths are those below the first quartiles of the median tail length, “median” lengths fall between the first and third quartiles, and “long” lengths are above the third quartile **(Figure 3C and Supplemental table 5)**. A gene is typically classified into one of the 3 groups. However, we observed that it is common for genes with differing isoforms to exhibit different polyA tail length, whereby ∼ 45% of genes have transcripts that are categorized in the three groups **(Figure 3D)**.

**Figure 3.**
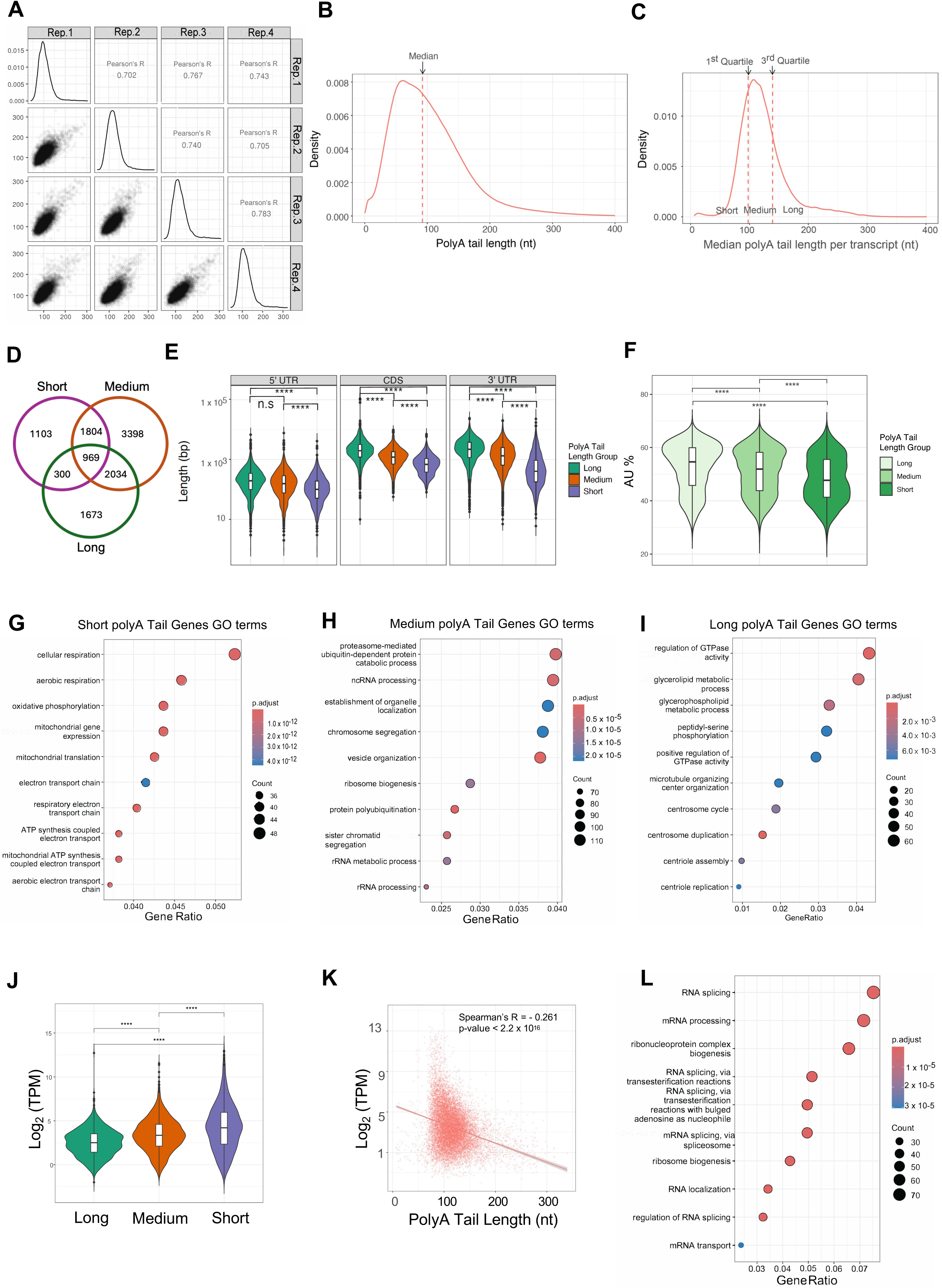
Genome wide characterization of poly(A) tail length and its correlation with mRNA abundance in leukemia cells. **(A)** Pairwise scatterplots of Pearson’s correlation showing the median polyA tail length per transcript across four control replicates. Each transcript has at least 10 reads. **(B)** Distribution of polyA tail length measured for each single read in control cells. Vertical lines represent the median polyA tail length of 92 nucleotides. **(C)** The distribution of median polyA tail lengths measured for each transcript when combined all reads in control cells. Each transcript has at least 10 reads. **(D)** Venn diagram showing overlapping gene names under each polyA tail length group in control cells (long vs. medium vs. short). **(E)** The length of RNA features (5’UTR, CDS, and 3’UTR) of transcripts categorized into each polyA tail length group (long vs. medium vs. short). Wilcoxon test, * p < 0.05, ** p < 0.005, *** p < 0.0005, **** p < 0.00005. **(F)** Assessment of AT content of transcripts categorized into each polyA tail length group (long vs. medium vs. short). Wilcoxon test, * p < 0.05, ** p < 0.005, *** p < 0.0005, **** p < 0.00005. **(G-I)** GO analysis of genes categorized into each polyA tail length group i.e. **(G)** “short”, **(H)** “medium”, **(I)** “long”. p-values are adjusted by Benjamini-Hochberg method with threshold p.value < 0.01 & q.value < 0.05. Top 10 pathways are shown. **(J)** TPM distribution of transcripts categorized into each polyA tail length group (long vs. medium vs. short). Wilcoxon test, * p < 0.05, ** p < 0.005, *** p < 0.0005, **** p < 0.00005. **(K)** Scatterplot showing correlation between averaged polyA tail length of individual transcripts vs. transcript expression (TPM) in control cells. Spearman’s R = -0.261, p. value < 2.2 x 10^-16^. **(L)** GO analysis of genes showing significant Spearman’s correlation in (K) Spearman’s R|≥ 0.1 & p. value < 0.005. p-values are adjusted by Benjamini-Hochberg method with threshold p. value < 0.05 & q. value < 0.2. Top 10 pathways are shown.

To look for potential RNA features influencing polyA tail lengths, we examined the lengths of various RNA features including CDS, 5’UTR, and 3’UTR and found that longer features exhibit significantly longer polyA tails **(Figure 3E)**. In addition, we saw that the sequence composition, in particular the rich content of AU (% AU) displayed positive correlation with polyA tail lengths **(Figure 3F)**. To examine whether polyA tail lengths can represent a defined characteristics of genes involved in particular cellular activities, we performed Gene Ontology analysis for genes found uniquely in the three short, medium and long groups. We found that genes with short polyA tails are highly enriched in pathways related to cellular respiratory and mitochondrial activities **(Figure 3G)**; genes with medium polyA tails are involved in protein catabolic process and rRNA biogenesis (**Figure 3H**) and genes with long polyA tails are dominantly represented in regulation of GTPase activity, metabolism of glycerolipids and centrosome (**Figure 3I**). These observations pointed to the diversity of polyA tail lengths associated with RNA features and gene functions, suggesting involvement of control of polyA tail lengths in directing molecular and cellular processes.

Given the general concept of positive correlation between polyA tail length and mRNA stability and expression level, we examined the transcript expression levels within the three groups categorized by polyA tail lengths. We observed a significant increase in transcript expression levels from “long” to “medium” and “short” groups **(Figure 3J).** Global analysis revealed a considerable negative correlation between gene expression and median poly A tail lengths (Spearman’s R = -0.26, p-value = 2.2 x10^16^), with highly expressed genes typically having shorter polyA tails **(Figure 3K)**. While the results challenged the dogmatic assumption that long polyA tailed transcripts are better protected from degradation, hence often express at higher levels; our observation of an inverse relationship between polyA tail length and transcript expression is consistent with findings from previous reports^37, 40^. Interestingly, we noticed that a large portion of genes exhibiting robust negative correlations (Biological Pathway, Benjamini-Hochberg p-value < 0.05) belong to pathways associated with RNA splicing events **(Figure 3L)**, suggesting a potential modulation of RNA splicing activity by mechanisms controlling polyA tail lengths.

### Correlation analysis of m^6^A RNA modifications and polyA tail length

One of the well characterized consequences of m^6^A modifications is mRNA degradation mediated by the CCR4-NOT complex recruited by the m^6^A YTHDF2 reader protein^13^. Subsequently, the CCR4-NOT complex deadenylates the polyA tail, leading to shortening of the tail and destabilization of the m^6^A marked transcripts. These results extrapolated a direct correlation between m^6^A modification and the length of the polyA tail. However, no direct assessment was performed to test the hypothesis due to the lack of simultaneous evaluation of m^6^A and polyA tail on the same datasets. Our evaluation of both m^6^A and polyA tail not only on the same cells but on the same mRNA molecule gave us a unique opportunity to address this question. We investigated the association between the mean m^6^A modification ratio and the length of the polyA tail at the transcript level and only observed a very weak global correlation (Spearman’s R = 0.038, p-value = 0.015) **(Figure 4A)**. Next, we looked at the distribution of polyA tail length in 5 subgroups of transcripts with increasing m^6^A modification ratio and also found no significant correlation **(Figure 4B)**. We further stratified transcripts based on the number of m^6^A sites and observed a small, significant increase in median polyA tail length in groups of transcripts with more m^6^A sites **(Figure 4C)**. In addition, we looked at the potential correlation between modified (m6Anet probability ≥ 0.6) vs. non-modified (m6Anet probability = 0) transcripts of a same gene and found only a few genes showing statistically significant alternations (Mann-Whitney test, p-value < 0.05, |estimate| ≥ 10) (**Figure S5**). These data indicate that m^6^A modifications are not negatively correlated with genome wide polyA tail length at the steady state.

**Figure 4.**
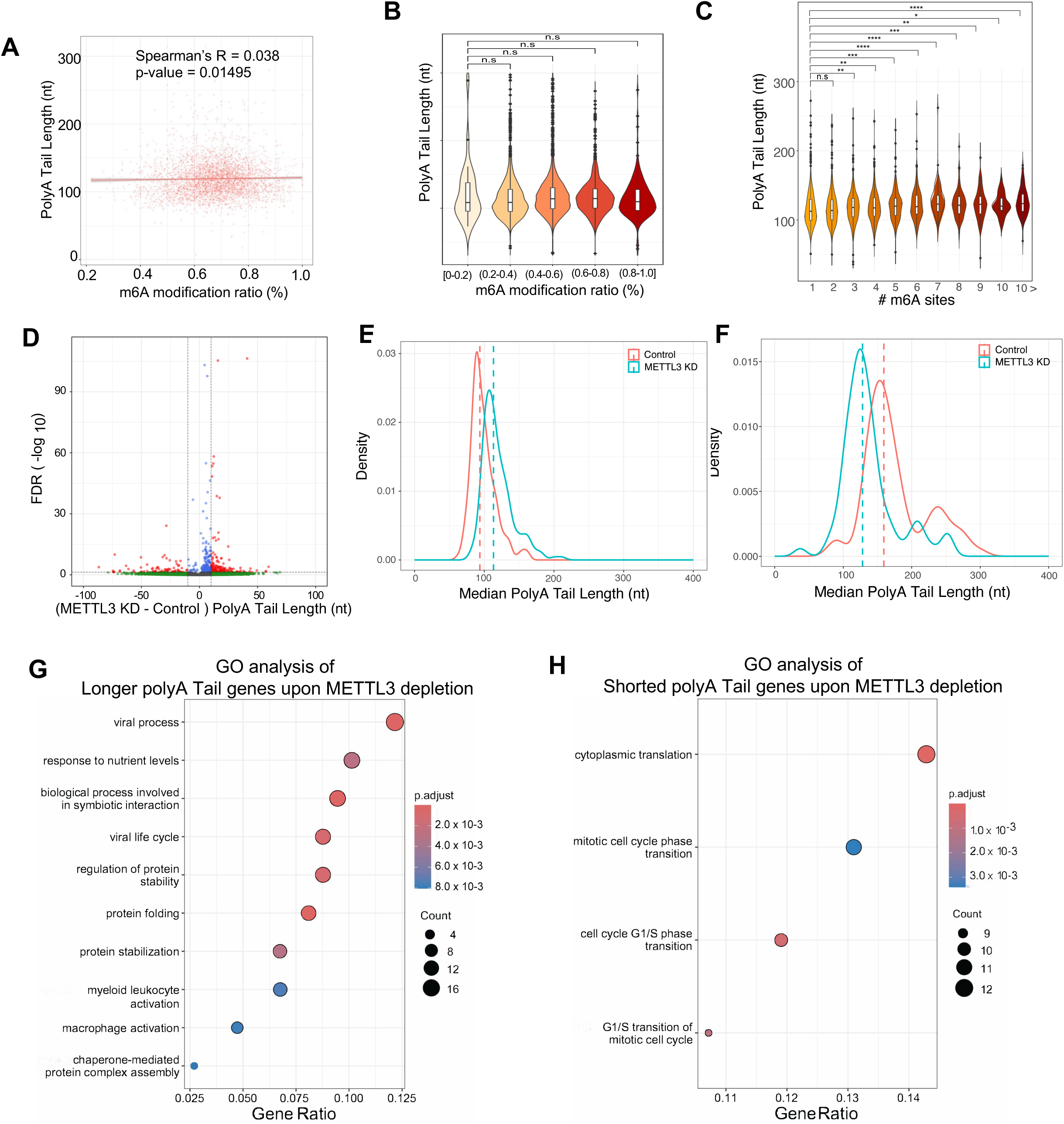
Correlation analysis of m^6^A RNA modifications and polyA tail length. **(A)** Scatterplot showing correlation between averaged m^6^A modification ratio matched with averaged polyA tail length of each transcript in control cells. Spearman’s R = 0.038, p-value = 0.01495. **(B)** The distribution of median polyA tail length of transcripts categorized based on the average m^6^A modification ratios (m6Anet probability ≥ 0.7) **(C)** The distribution of median polyA tail length of transcripts categorized based on number of m^6^A sites on gene level (m6Anet probability ≥ 0.9). **(D)** Volcano plot of Mann-Whitney test results comparing changes in polyA tail length of individual transcripts in METTL3 depleted cells (METTL3 KD) vs. control (|KD - control| = 10 (nt), FDR < 0.05). **(E)** Density plot of median polyA tail length of group of transcripts when polyA tail length were decreased upon METTL3 depletion. **(F)** Density plot of median polyA tail length of group of transcripts when polyA tails were increased upon METTL3 depletion. **(G-H)** GO analysis of genes showing (**G**) longer tails upon METTL3 depletion and (**H**) shorter tails upon METTL3 depletion. p-values were adjusted using the Benjamini-Hochberg method with a threshold of p-value < 0.01 & q-value < 0.05. The top 10 pathways are shown.

Next, we wanted to examine whether dynamic changes in m^6^A can result in alternation of polyA tail length at the transcript level. We performed statistical analysis employing the Mann-Whitney test (read count > 30; |Δ PolyA Tail Length| > 10 and FDR < 0.05) to define transcripts showing significant changes in median polyA tail length upon METTL3 depletion (**Figure 4D and Supplemental table 5**). We observed a larger number of transcripts showing increased polyA tail length in METTL3 KD cells. 186 transcripts exhibit a significant increase in polyA tail length (93.7 vs. 113 nucleotides polyA tail lengths for control vs. METTL3 KD -**Figure 4E**), whereas 100 transcripts exhibited significantly shorter polyA tails upon METTL3 depletion (159 and 128 nucleotides for control vs. METTL3 KD-**Figure 4F**). GO term analysis (Biological Pathway, Benjamini-Hochberg p-value < 0.01) demonstrated that genes with longer polyA tail lengths after METTL3KD are enriched in pathways related to viral process, myeloid leukocyte activation and protein folding (**Figure 4G**) while genes exhibiting decreased polyA tail length after METTL3KD are involved in cell cycle regulation (**Figure 4H**). The results suggest that while the correlation is not global, regulation of m^6^A and polyA are possibly coordinated for a subset of m^6^A targets under METTL3 control.

### METTL3-mediated m^6^A regulation and alternative splicing control

m^6^A modifications have been showed to influence RNA splicing^41^. The presence of m6A at the splice site can hinder binding of splicing factors, thereby preventing processing of pre-mRNAs^41^. Dysregulation of METTL3-mediated m^6^A modifications can result in aberrant expression of splicing factors, contributing to transformation and cancer phenotypes^42^. Given the functional interaction between m^6^A RNA methylation and splicing, we explored the correlation between the presence of m^6^A and splicing events and how alternations in m^6^A upon METTL3 depletion influence alternative splicing in leukemia cells. We used Isoquant^43^ to quantify isoforms from dRNA-seq data. For the analysis of alternative splicing, we used FLAIR^44^ to identify differential alternative splicing events between control samples and those with METTL3 knockdown.

To identify alternative splicing events altered in METTL3 depletion, we conducted a differential splicing event analysis using Fisher’s exact test via FLAIR. We identified four categories of alternative splicing events, including alternative 3’ splice site (A3’SS), alternative 5’ splice site (A5’SS), cassette exon (ES), and intron retention (IR), encompassing a total of 233-4217 splicing events within each category **(Figure 5A and Supplemental table 6)**. The most prominent changes were observed in cassette exon and intron retention. The alternative splicing events mapped to 45 to 599 isoforms and 16 to 121 genes **(Figure 5B-C)**. A Venn diagram illustrating the overlap between the four alternative splicing categories at the gene level showed little overlap. This suggests that the majority of splicing events are unique for individual genes **(Figure 5D)**. GO analysis of 171 genes showing differential alternative splicing between control and KD cells captured biological processes of cytoplasmic translation and metabolic pathways involved in the biosynthesis and regulation of nucleotides, including purine ribonucleoside triphosphates and ATP (**Figure 5E and Figure S6A-B**).

**Figure 5.**
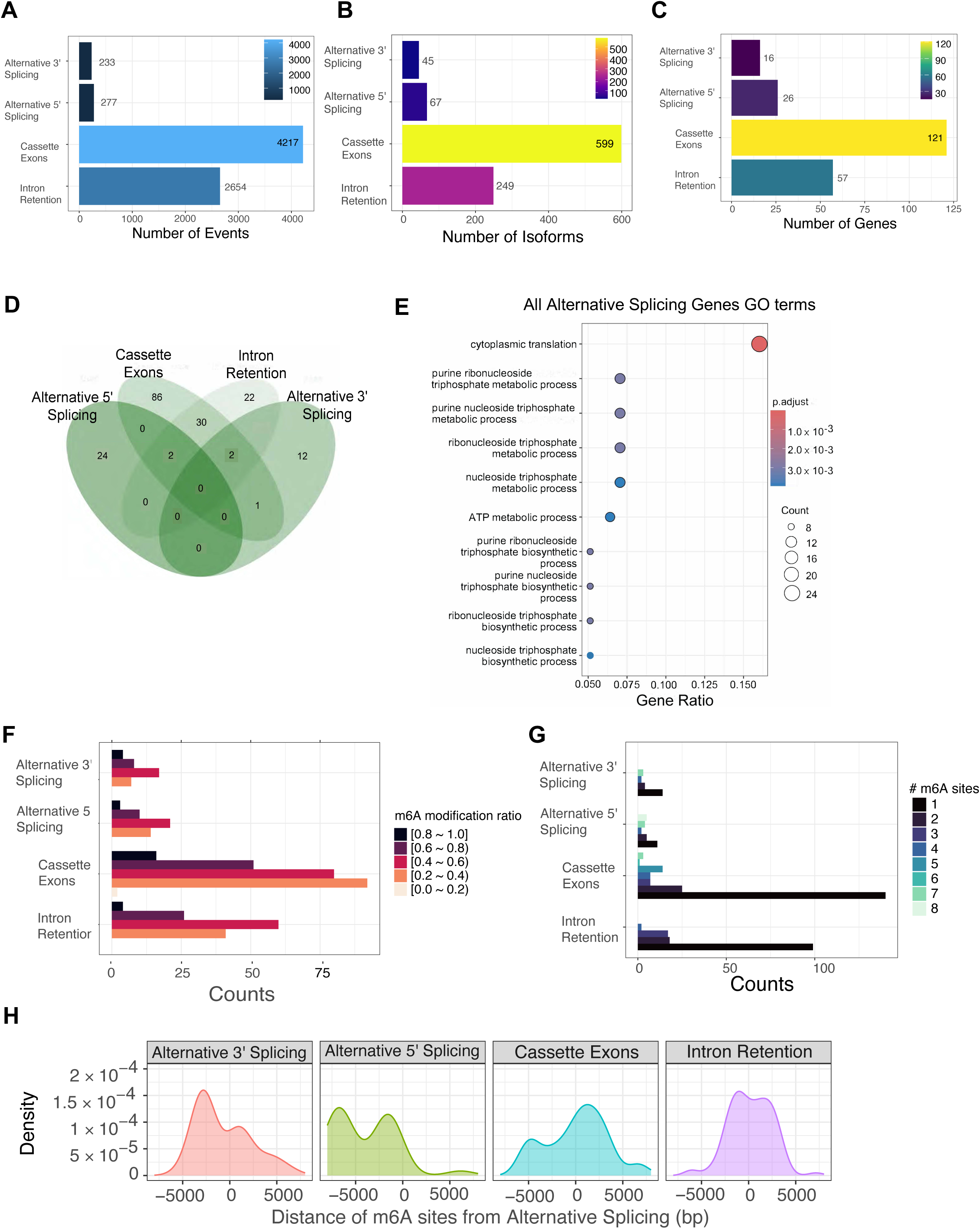
METTL3-mediated m^6^A regulation and alternative splicing control. **(A)** Bar plots showing event counts for alternative RNA splicing identified in METTL3 depleted (METTL3 KD) vs. control cells. Fisher’s Exact Test, p-value < 0.05. 4 main types of alternative splicing were characterized i.e., alternative 3’ splicing; alternative 5’ splicing; Cassette Exon and Intron retention. **(B)** Bar plots showing isoform counts for alternative RNA splicing events in METTL3 depleted (METTL3 KD) vs. control cells. Fisher’s Exact Test, p-value < 0.05. **(C)** Bar plot of gene counts for alternative RNA splicing events in METTL3 depleted (METTL3 KD) vs. control cells. Fisher’s Exact Test, p-value < 0.05. **(D)** Venn diagram showing overlapping genes found to have different alternative splicing patterns. **(E)** GO analysis result of gene list showing differential alternative splicing events between METTL3 depleted (METTL3 KD) vs. control cells. p-values are adjusted by Benjamini-Hochberg method with threshold p. value < 0.05 & q. value < 0.2. Top 10 pathways are shown. **(F)** Bar plots showing m^6^A modification ratio (m6Anet probability ≥ 0.7) by isoforms categorized into 4 types of alternative splicing. **(H)** Bar plots showing number of m^6^A modification sites by genes categorized into 4 types of alternative splicing. **(I)** Density plots showing distance of m^6^A modification sites from the alternative splicing events.

To examine potential associations between alternative splicing events and m^6^A modifications, we assessed the m^6^A modification ratio (**Figure 5F**) as well as number of m^6^A sites (**Figure 5G**) of each spliced transcript categorized into the 4 groups. Interestingly, we observed that isoforms whose RNA methylation frequencies range from 0.2-0.4 and 0.4-0.6 (20-60% of mRNA molecules is modified) had the highest rate of changes in splicing. Despite the fact that there are more transcripts with higher modification ratio (0.8-1.0) (**Figure 1G**), fewer alternative splicing events were found in this group (**Figure 5F**). At the same time, more events were also mapped to isoforms with only 1 or 2 m^6^A sites (**Figure 5G**). Notably, cassette exon group is the one showing different profiles than the other 3 groups. These observations suggest potential differential regulation of the type of splicing control by m^6^A modifications. We then asked whether the location of the m^6^A sites has any effect on the splicing events. We saw distinct patterns of the spatial proximity of m^6^A sites to the sites of splicing events across the 4 groups (**Figure 5H**). Taken together, these results demonstrate that long-read dRNA-seq can unveil patterns of m^6^A associated with specific alternative splicing events, offering additional insights into how the two regulatory mechanisms potentially coordinate to regulate RNA processing.

## Discussion

In this study, we demonstrated the use of dRNA-seq to extensively investigate the transcriptome and epitranscriptome of human leukemia cells. We obtained the sufficient sequencing coverage and depth necessary to conduct an unbiased and genome wide analysis. We showed that the approach effectively captured many features of an mRNA molecule including gene and transcript abundance, precise mapping of m^6^A modifications, quantitative estimation of m^6^A modification rates and measurements of polyA tail length distribution genome wide as well as at the gene and transcript levels, and characterization of RNA isoforms and differential alternative splicing events. Notably, as these features were acquired simultaneously within the same samples, it was possible to perform correlative analysis to directly assess potential interactions between multiple post-transcriptional modifications within individual RNA molecules. Indeed, we provide strong evidence to validate previous reports^40^ on the negative correlation between polyA tail length and transcripts’ abundance at a steady state. Short polyA tails are more frequently associated with abundant transcripts and genes. In addition, we did not find a strong correlation between m^6^A modifications and polyA tails. These data support the revision of our current view of how polyA tails function to regulate gene expression^45^. Overall, our high-quality datasets enabled unprecedent exploration and offered a valuable resource for the field as the rich dRNA-seq outputs can be further exploited to extract more information and insights, such as information on additional RNA modifications^46^.

Through our investigation, we also uncovered a more nuanced regulatory mode of m^6^A on mRNA transcripts. By assessing m^6^A stoichiometry alongside gene expression changes, we demonstrate that while m^6^A does not have a strong effect on steady state expression of mRNA, variation in the number of m^6^A sites and m^6^A stoichiometry on an mRNA substrate may indeed regulate its expression. Of course, this raises additional questions. For example, if an mRNA transcript contains multiple m^6^A modifications, does the presence of multiple m^6^A decorations enable YTHDF2 binding, and as a consequence, a higher likelihood of mRNA degradation? Moreover, it has previously been suggested that m^6^A modifications in mRNAs increase their respective translational rate, whereby YTHDF1 binds to m^6^A moieties and mediates the interaction of mRNAs with the eIF3 translation initiation factor^47^. However, little is still known about the influence multiple m^6^A decorations along an mRNA may play in regulating translational processes. Furthermore, we demonstrate that the METTL3 derived regulation of m^6^A sites in mRNAs appears to be pathway specific. In many cases, hypermethylation and hypomethylation both correspond with down and upregulated transcripts when METTL3 is knocked down. As such, our study demonstrates that a rich dRNA-seq dataset is invaluable for the analysis of RNA modifications like m^6^A and as the technology improves and more features may be extracted from these types of data, the value of datasets like these will only improve.

The efficacy of dRNA-seq is determined by several factors including base-calling accuracy, signal segmentation, read-depth, and mapping of elongated sequencing reads. Improvement in any of those parameters can influence the quality of dRNA-seq outputs. Typically, a 30X coverage is used to select for unique transcripts^23^. Given the increased accuracy and the fact that actual reads are mapped directly to a transcript, a 20X^33^ or 10X^28^ coverage is also employed as an applicable threshold. To identify m^6^A sites, we used m6Anet^33^, a software based on a neural-network-driven multiple-instance learning (MIL) model. The major advantage of m6Anet is the de novo identification of the modification without a need for a paired comparison against unmodified samples as in comparative methods such as Nanocompore^23^, xPore^48^, and ELIGOS^49^. However, the method does favor detection of the modification sites to the consensus motif DRACH of m^6^A, hence likely misses out the mapping of non-DRACH sites. There are a number of additional methods such as TOMBO^50^, Yanocomp,^51^ Epinano^52^ and more recently, CHEUI^53^ which have been developed to measure m^6^A modifications in dRNA-seq datasets. With the development and optimization of computational tools and the arrival of the new RNA chemistry for dRNA-seq kit, it is expected that further improvement and wider adoption of dRNA-seq for study of RNA biology will be achieved in the future.

## Supplemental Figure legends

**Figure S1.**
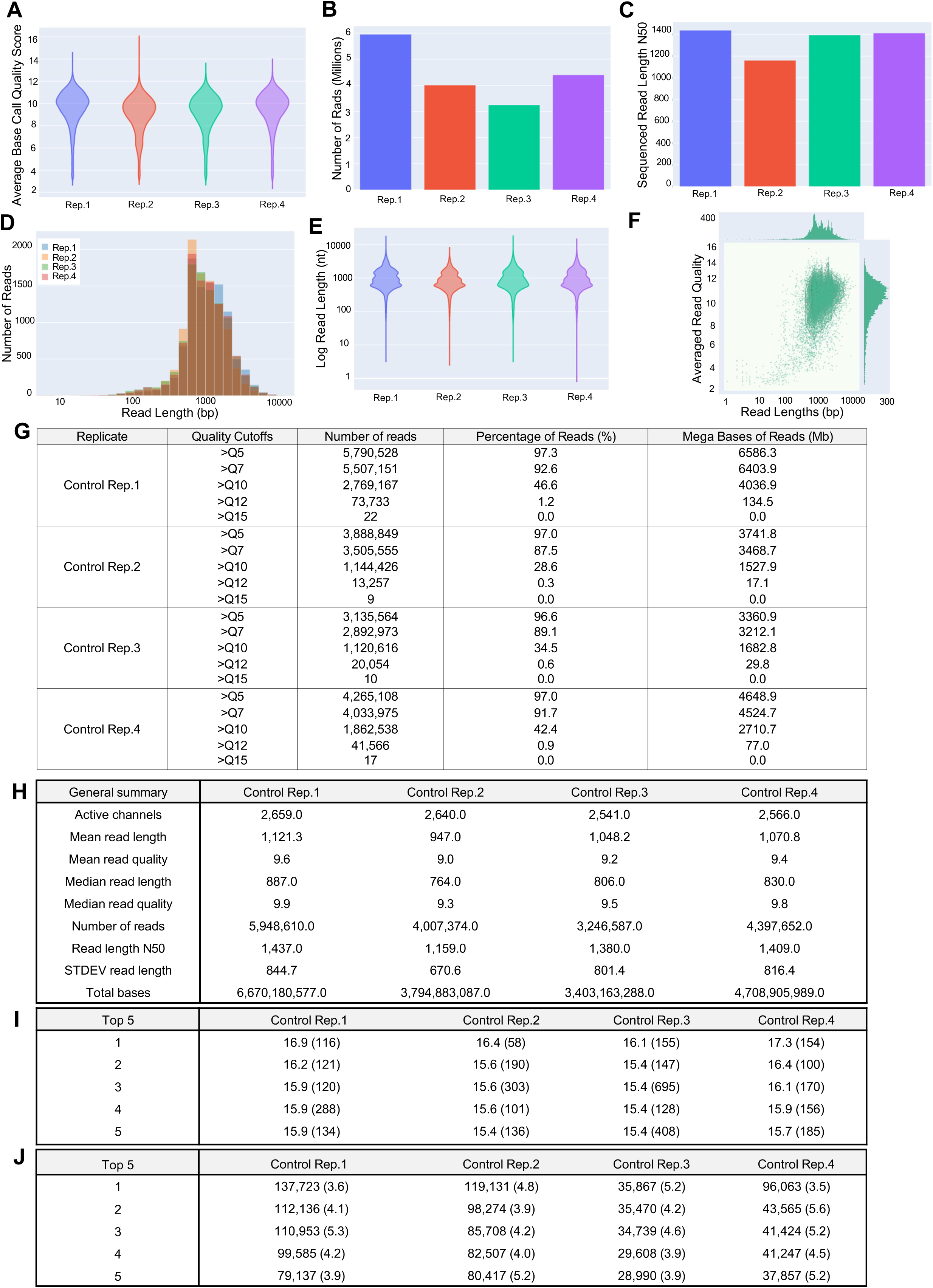
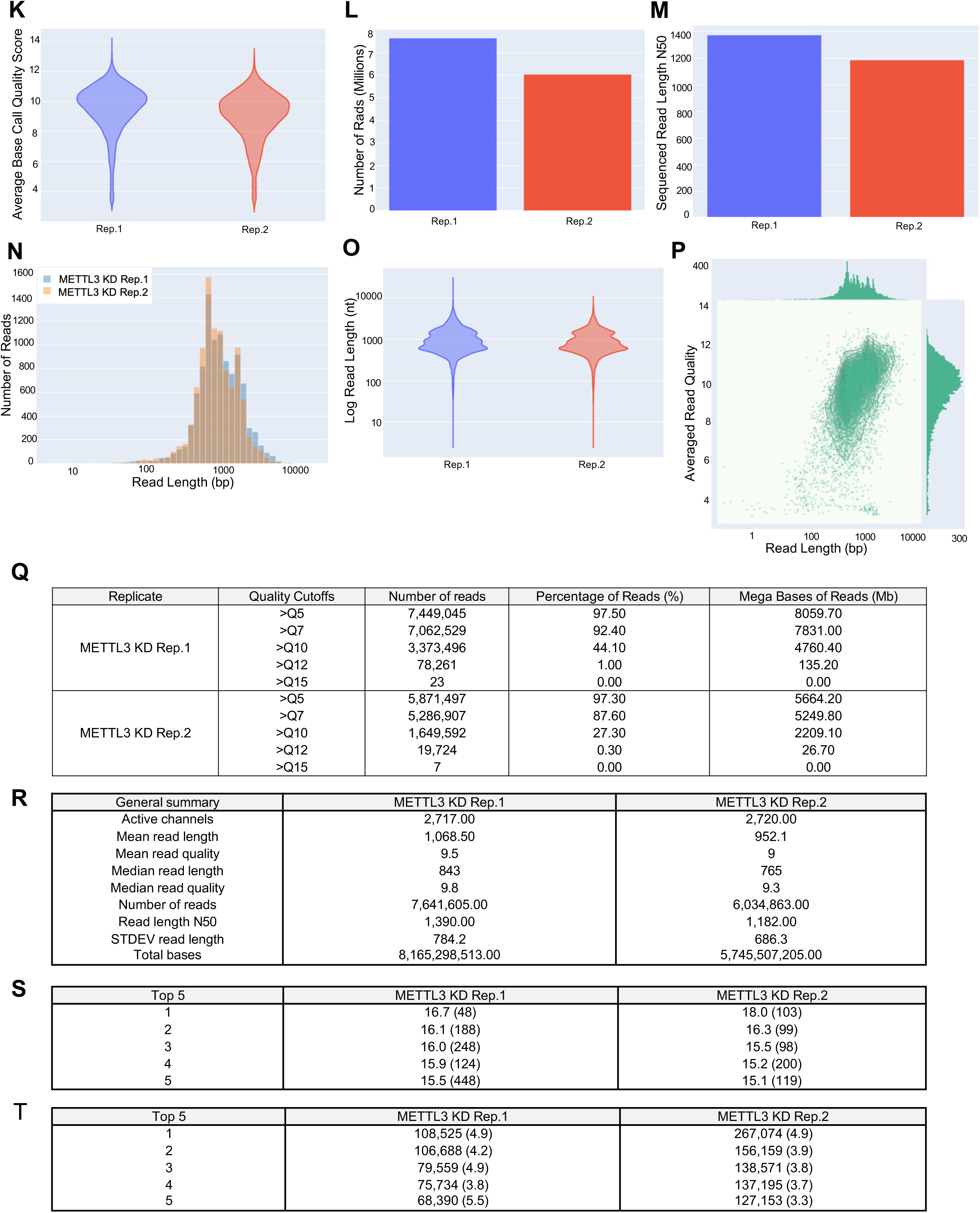
**(A)** Violin plots depicting the average base call quality score in 4 control replicates. **(B)** Bar plots showing number of reads in 4 control replicates. **(C)** Bar plots showing read length N50 in 4 control replicates. **(D)** Histogram of log transformed read lengths in 4 control replicates. **(E)** Violin plots showing log-transformed read length in 4 control replicates. **(F)** Bivariate plot comparing log transformed read length with average basecall Phred quality score (showing representative data from control Replicate 3) **(G)** Table displaying the number, percentage, and mega bases of reads above quality cutoffs in 4 control replicates. **(H)** Summary table of dRNA seq runs in 4 control replicates. **(I)** Top 5 highest mean base call quality scores in control replicates (with read lengths)**. (J)** Top 5 longest reads in control replicates (with mean base call quality score)**. (K)** Violin plots depicting average base call quality score in 2 METTL3 knocked down (METTL3 KD) replicates. **(L)** Bar plots showing number of reads in METTL KD replicates. **(M)** Bar plots showing read length N50 in METTL KD replicates. **(N)** Histogram of log transformed read lengths in METTL KD replicates. **(O)** Violin plots of log-transformed read length in METTL KD replicates. **(P)** Bivariate plot comparing log transformed read length with average basecall Phred quality score ((showing representative data from METTL KD replicate 1). **(Q)** Table displaying the number, percentage and mega bases of reads above quality cutoffs in KD replicates. **(R)** Summary table of dRNA seq runs in METTL KD replicates. **(S)** Top 5 highest mean base call quality scores in KD replicates (with read lengths). **(T)** Top 5 longest reads in KD replicates (with mean base call quality score).

**Figure S2.**
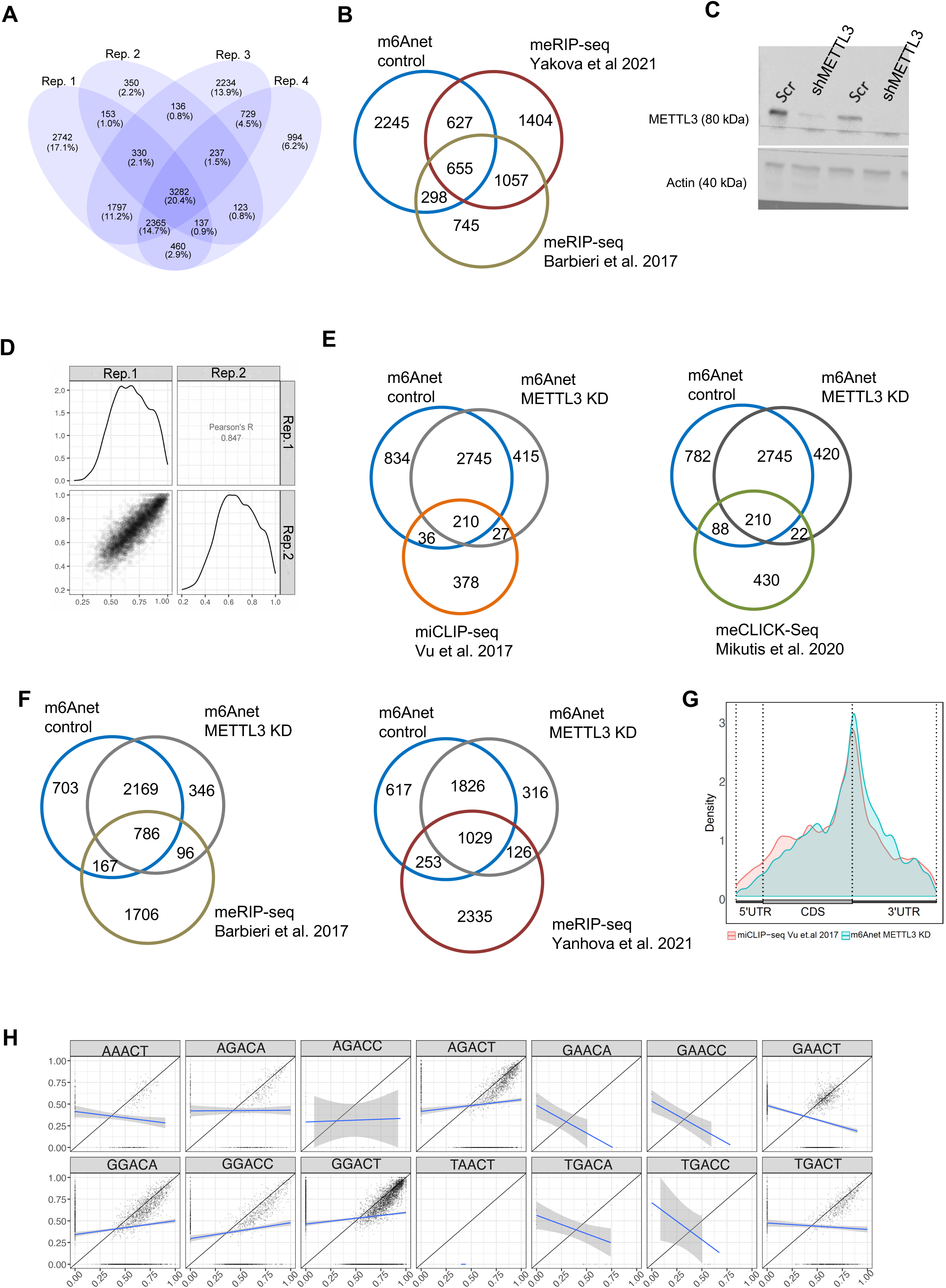
(A) Venn diagram showing overlapping m^6^A genes identified by dRNA-seq in 4 control replicates. (B) Metagene plot depicting the transcriptome-wide distribution of m^6^A across 5’-untranslated regions (UTRs), coding sequences (CDS), and 3’-UTRs from nanopore direct RNA-seq (m6Anet probability ≥ 0.9) and miCLIP-seq (Mikutis et al. 2020) respectively. (**C**) Immunoblots demonstrating efficient depletion of METTL3 in MOLM13 cells transduced with shRNA against METTL3 (METTL3 KD) vs. scramble control shRNA (control). ACTIN serves as loading control. (**D**) Pairwise scatterplots of Pearson’s correlation of m^6^A modification ratios for sites common (m6Anet probability ≥ 0.9) across two METTL3 replicates (Pearson’s R = 0.85, p-value < 0.05). The number of reads for each transcript > 10. (**E-F**) Venn diagram showing overlapping m^6^A genes identified in dRNA-seq (m6Anet probability ≥ 0.9) of control and METTL3 KD cells and previously identified m^6^A targets by other m^6^A profiling method. (**G**) Metagene plot illustrating the transcriptome-wide distribution of m^6^A across 5’-untranslated regions (UTRs), coding sequences (CDS), and 3’-UTRs from dRNA-seq of METTL3 KD samples (m6Anet probability ≥ 0.9) and miCLIP-seq (Vu et al. 2017) respectively. **(H)** Scatterplots showing comparison of m6Anet m^6^A modification ratios (m6Anet probability ≥ 0.9) between control and METTL3_KD cells based on DRACH motifs.

**Figure S3.**
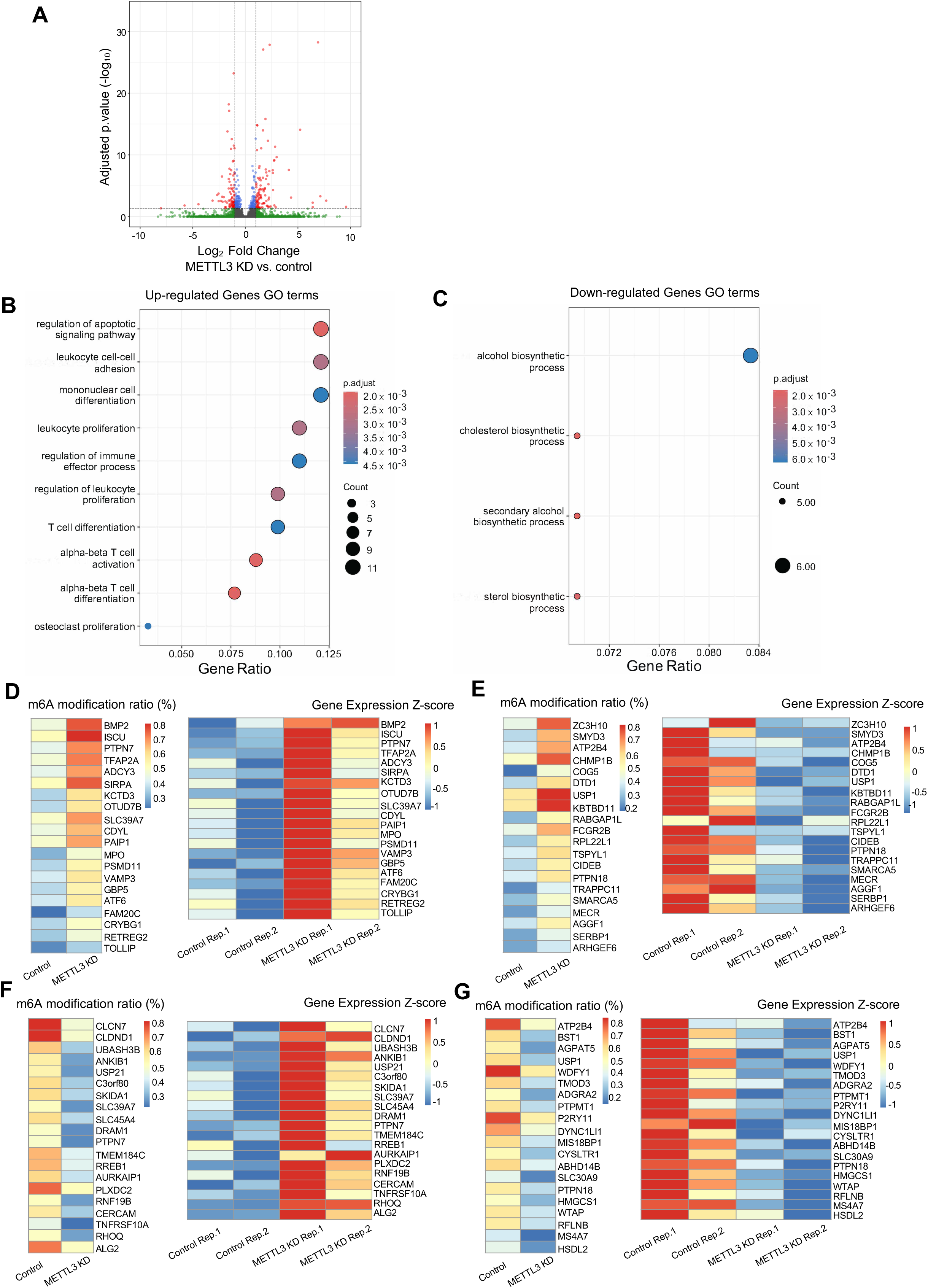
**(A)** Volcano plot showing differential gene expression (DEG) analysis of METTL3_KD vs. control. (|Log_2_ Fold Change| ≥ 1, adjusted p. value < 0.05). **(B-C)** GO analysis of gene groups (**B**) upregulated Log_2_ Fold Change ≥ 1 (adjusted p.value < 0.05) and (**C**) downregulated Log_2_ Fold Change ≤ -1 (adjusted p.value < 0.05) upon METTL3 depletion. p-values are adjusted by Benjamini-Hochberg method with threshold p. value < 0.01 & q. value < 0.05. **(D-G)** Heatmaps presenting significant conjoint analysis results of m^6^A methylation levels and gene expression (CPM) in Figure 2E: (**D**) Hyper-methylated & Up-regulated (hyper-up), (**E**)Hyper-methylated & Down-regulated (hyper-down), (**F**) Hypo-methylated & Up-regulated (hypo-up) and (**G**) Hypo-methylated & Down-regulated (hypo-down).

**Figure S4.**
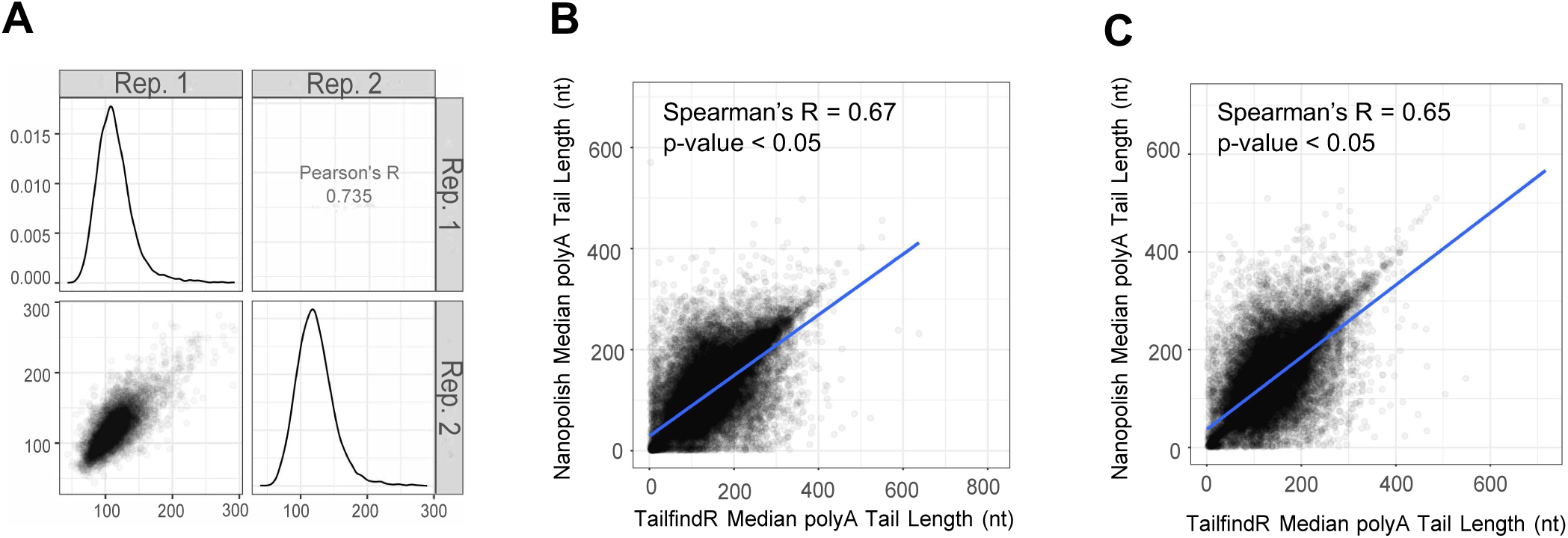
**(A)** Pairwise scatterplots showing the median poly A tail length per transcript across two METTL3 KD replicates. The number of reads for each transcript > 10. **(B-C)** Scatterplots depict median polyA tail length per transcript from tailfindR and Nanopolish **(B)** in control cells, Spearman’s R = 0.67, p-value < 0.05 and **(C)** METTL3 KD cells, Spearman’s R = 0.65, p-value < 0.05.

**Figure S5.**
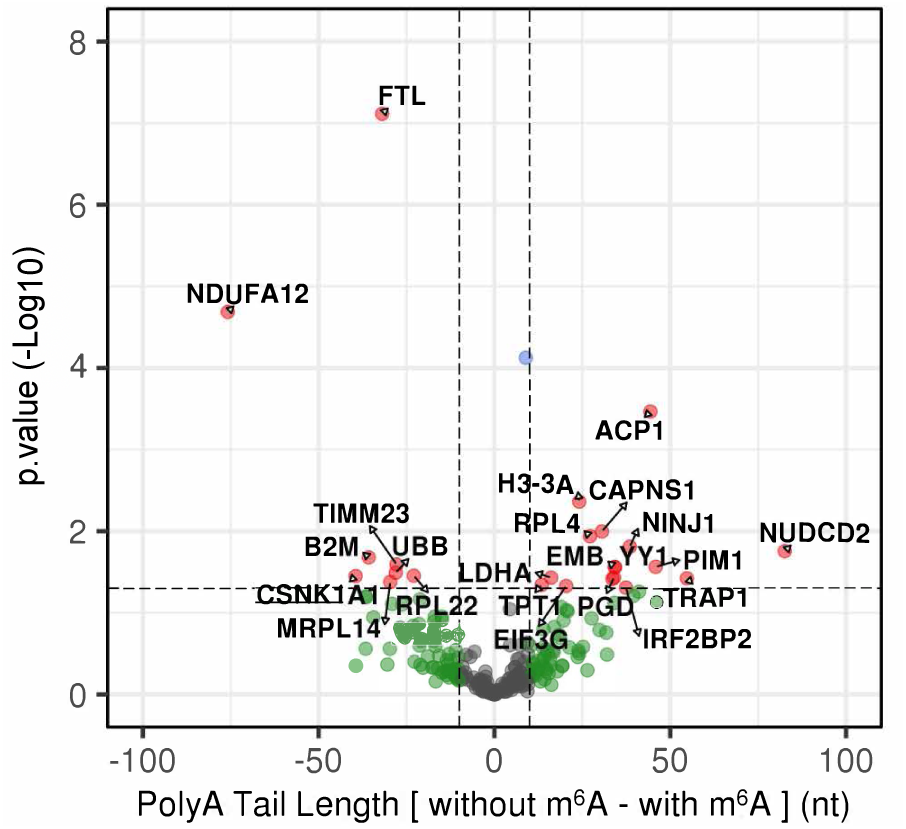
Volcano plot showing Mann-Whitney test results comparing polyA tail length on the gene level between two groups, ‘without m^6^A’ (m6Anet probability = 0) and ‘with m^6^A’ (m6Anet probability ≥ 0.6) (|without m^6^A – with m^6^A | ≥ 10, p.value < 0.05) in control cells. Transcripts with a read count ≥ 10 from each group were subsetted.

**Figure S6.**
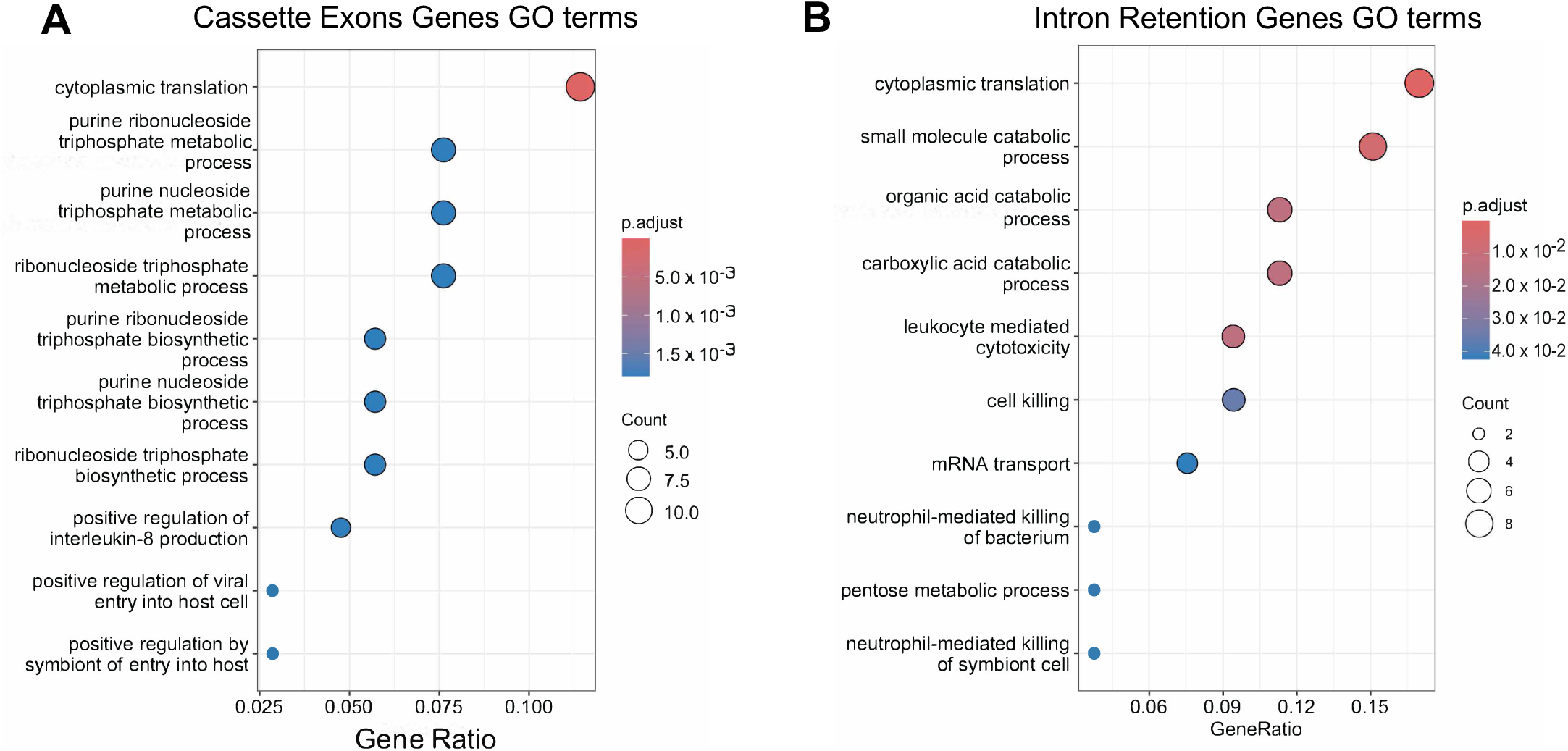
**(A-B)** GO analysis of genes with significant alternative splicing of **(A)** Cassette exons, 114 unique gene names in total. **(B)** Intron retention, 55 unique gene names in total. p-values are adjusted by Benjamini-Hochberg method with threshold p. value < 0.05 & q. value < 0.2. Top 10 pathways are shown.

## Methods

### Tissue culture

Human myeloid leukemia cell line MOLM13 (classified under AML French-American-British (FAB) M5a) was cultured in RPMI-1640 medium supplemented with 10% fetal bovine serum (FBS), penicillin (100 units/ml), and streptomycin (100 units/ml). Adherent cell line HEK293T was cultured in DMEM medium supplemented with 10% FBS, penicillin (100 units/ml), and streptomycin (100 units/ml). These cell cultures were maintained at 37°C in a 5% CO2 humidified incubator. The cell lines utilized in this study were procured from the American Type Culture Collection (ATCC) and rigorously screened to confirm their absence of mycoplasma contamination.

### Lentivirus production and transduction

pLKO.1 lentiviral plasmids carrying scramble shRNA control and shRNA targeting METTL3 were previously validated^8^. HEK293T cells were cultured in 10cm plates until reaching 90% confluence prior to cell transfection. The lentiviral transfer vector DNA, in conjunction with psPAX2 packaging and pMD2.G envelope plasmid DNA, were combined in a specific ratio of 4:3:1 (12.5 μg : 9.375 μg : 3.125 μg) in 1 mL of 0.25M CaCl2. This mixture of plasmid DNA was cautiously introduced dropwise into 1 mL of filtered 2x concentrate BES buffered saline (Millipore 14280, bioWORLD 40220006-2), while gently inducing air bubbles through the DNA mixture using a sterile 1 mL pipette. After a 5-minute incubation period at room temperature, the solution was incrementally added to the HEK293T cells. Subsequently, the plates were delicately rocked in a circular motion to evenly distribute the precipitates before returning them to a 5% CO2 incubator set at 37°C under humidified conditions. Following a 16-hour incubation period, the media was replaced with fresh culture medium. Viral supernatants were harvested twice, at 24-hour and 48-hour intervals post-medium replacement. The supernatant underwent filtration through a 0.45 μm pore PVDF Millex-HV filter (Millipore, SLHVR33RB) and was stored at -80°C for subsequent use.

To transduce leukemia cells with viral particles, MOLM13 cells underwent transduction using lentiviruses via spinfection at a speed of 1400 revolutions per minute (rpm) for a duration of 1 hour at room temperature. Subsequently, the cells were seeded in RPMI supplemented with 10% fetal bovine serum (FBS) at a density of 0.5x106 cells/ml. To enable puromycin selection, a concentration of 3 μg/ml puromycin was introduced to the transduced cells at 24 hours post-transduction. After an approximate period of 48-56 hours post-transduction, during which a puromycin selection phase spanning approximately 24-32 hours was implemented, cells were harvested. The selected cells were then gathered for subsequent Western blot analyses and nanopore direct RNA sequencing procedures.

### Immunoblotting

Cells were quantified and subjected to two washes with cold phosphate-buffered saline (PBS) before being collected. A total of 200,000 cells were lysed in 40 μL of 1X Laemmli protein loading buffer and subsequently boiled at 95°C for 5 minutes. The resultant whole-cell lysates were separated using either 4%–15% TGX precast protein gels or 4–15% Criterion™ TGX™ Precast Midi Protein Gel (BioRad, 4561085, 5671083, 5671084) and then transferred onto nitrocellulose membranes (BioRad, 1620115). These membranes were blocked with 5% milk for 1 hour before being subjected to antibody incubation against specific targets, including METTL3 (Thermo Fisher, 150731-1-AP, 1:1000) and ACTIN (Sigma Aldrich, A3854, 1:5000). Following an overnight incubation with primary antibodies, the membranes underwent washing with 1 × PBST and subsequent incubation with HRP-linked secondary goat anti-rabbit IgG [NEB, 7074V], 1:3000) for 1 hour at room temperature. Protein bands were visualized using Immobilon ECL Western HRP Substrate (Millipore, WBKLS0500) and ECL Western Blotting Detection Reagent (Sigma, GERPN2105) via the Bio-Rad ChemiDoc Imaging System employing chemiluminescence detection. The relative expression levels of each protein band were quantitatively analyzed using Image J software, with β-actin serving as the internal reference standard.

### Direct RNA sequencing

RNA was extracted from cell pellets using the vendor recommended protocol for trizol RNA extraction. Briefly, trizol [Invitrogen, 15596026] homogenized cells were mixed with chloroform (0.2mL / 1mL trizol) for three minutes and centrifuged at 12,000 × g at 4°C for 15min. The aqueous phase was transferred to a new tube and mixed with isopropanol (0.5mL per 1mL trizol) for 10 minutes at 4°C and centrifuged at 12,000 × g at 4°C for 10min. The supernatant was discarded, and the RNA pellet was washed with 75% ethanol (1mL / 1mL trizol), centrifuged for 5 minutes at 7500 × g at 4°C, where the supernatant was subsequently also discarded. The RNA pellet was resuspended in pure RNase free water and 10-50 μg of RNA was DNaseI digested following vendor recommendations [Invitrogen, 18068015]. Total RNAs were poly(A) enriched using the NEBNext Poly(A) mRNA Magnetic Isolation Module [NEB, E7490L]: total RNA was diluted to 75μL and mixed 1:1 with prepared NEBNext Magnetic Oligo d(T)25 Beads at 65°C for 5min, then 5min at room temperature. The mixture was pelleted on a magnet and the supernatant was discarded, followed up by resuspension in wash buffer and immediately after pelleting again on the magnet. The supernatant was discarded and the wash step was repeated. The pellet was resuspended in 75μL Tris, incubated at 80°C for 2min, then mixed 1:1 with RNA binding buffer. After 5min at room temperature, samples were pelleted again on a magnet and the supernatant was discarded. Pellet was washed as previously described, resuspended in 12μL of nuclease free water and incubated for 2min at 80°C and 2min at room temperature. Samples were pelleted on a magnet again and the supernatant was transferred to a new tube.

Direct RNA libraries were generated using the direct RNA sequencing kit [ONT, RNA-002]. 9 μL RNA was mixed with 3μL Quick Ligation Buffer [NEB, E6056], 0.5μL RNA CS – spike in, 1μL RTA adaptor and 1.5μL T4 DNA Ligase [NEB, E6056] for 10min at room temperature. Then, a first strand synthesis reaction was initiated using 2μL 10mM dNTPs [NEB, N0447], 8μL first-strand buffer and 2μL Maxima H-Minus reverse transcriptase [Thermo, EP0752] in a final reaction volume of 40μL. The reaction incubated at 50°C for 60 min and 70°C for 10 min. The RNA:DNA hybrid was bead cleaned in a 1.8X reaction using the RNAClean XP bead system [Beckman, A63987] using manufacturer’s recommendations and eluting in 20μL nuclease free water. RMX adaptor ligation was performed using 8μL Quick Ligation Buffer, 6μL RMX adaptor, 3μL T4 DNA Ligase in a final reaction volume of 40μL and incubated for 15min at room temperature. The RMX ligated product was 1X bead cleaned as recommended by ONT: incubating the sample with RNAClean XP beads for 15min at room temperature. After pelleting on a magnet, the supernatant was discarded and the sample was resuspended in 150μL of ONT Wash buffer, pelleting immediately after. The wash step was repeated, and the pellet was resuspended in 41μL of ONT elution buffer, incubating for 10min at room temperature. After pelleting on a magnet, the supernatant was transferred to a new tube, where 40μL of sample was mixed with 75μL nuclease free water and 75μL ONT RRB reagent. The final prepared library was loaded onto an R9.4.1 PromethION flow cell on the PromethION 48 instrument following manufacturer’s recommendations. Sequencer operation was controlled by MinKNOW 22.08.6 using the direct RNA protocol. Basecalling was conducted by Guppy 6.2.7.

### dRNA-seq output preprocessing

Upon acquisition of raw signal and sequences from the PromethION 48 platform using the Guppy basecalling software^54^, the continuous signals undergo segmentation into discrete events associated with specific k-mers. Each occurrence of an event entails the collection of samples corresponding to the presence of a k-mer within the nanopore, followed by the subsequent event occurring as the nucleotide strand progresses through the pore by one base. The chosen k-mer length for RNA analysis is set at five.

Utilizing the ‘f5c eventalign’ approach^32^, signal fragments are paired with their corresponding reference nucleotides. Subsequent analysis involves the segmentation of each reading, delineating unique attributes inclusive of the reference k-mer, model k-mer events, associated signal segments, as well as the reported observed and expected normalized means. Multiple event means, weighted by their respective event length, are averaged to consolidate reported events occurring at the same position into a singular event. Skipped positions and mismatched k-mers were discarded from consideration. Positions within aligned reads ≥ 10 in length were deemed to offer sufficient coverage for subsequent analysis.

### Basecalling and alignment

The current signal of each FAST5 file was basecalled using Guppy v.6.5.7 and stored in FASTQ files. Minimap2 v.2.24^30, 31^ with the ‘-ax map-ont’ parameter was employed to align the reads to the transcriptome, utilizing GRCh38 Ensembl FASTA annotations from release version 106. Subsequently, only Ensembl transcript IDs that corresponded to reference annotations were retained for subsequent analyses.

### m^6^A modification ratio estimation

Estimation of the m^6^A modification ratio involved the use of m6Anet v.1.0^33^ to measure m^6^A sites by single-nucleotide resolution and m^6^A stoichiometry. Regions containing the DRACH motif were initially identified, and the m^6^A location, along with its modification probability and ratio, was subsequently determined. Regions with a coverage ≥ 10 were filtered and subjected to further analysis. For the comparison of significant changes in m^6^A levels at the single-nucleotide level between the control and KD cells, we utilized the Mann-Whitney test on modification probability values (|KD-control m^6^A probability| > 0.01, FDR < 0.05). Adjusted p-values were computed using the Benjamini–Hochberg method to control the false discovery rate (FDR). From miCLIP-seq datasets, m^6^A mapping results were extracted from mRNAs categorized as protein-coding genes and encompassing gene regions such as 5’ UTR, exon, and 3’ UTR. The m^6^A profiling results from Vu et al., 2017^8^ and Mikutis et al., 2020^29^ were downloaded and used for overlapping analysis. From meRIP-seq datasets, m^6^A mapping results were extracted from mRNAs belonging to protein-coding genes and covering gene regions including 5’ UTR, exon, and 3’ UTR. meRIP-seq datasets are Barbieri et al., 2017^18^ and Yankova et al., 2021^34^. All datasets were obtained from the MOLM13 cell line.

### Metagene analysis

Metagene analysis was conducted wherein gene coordinates were aligned to transcript coordinates utilizing ‘bedparse tx2genome’. Subsequently, m^6^A modification ratios per single nucleotide, possessing a high probability threshold ≥ 0.9, were selected as a subset. This subset was further used to superimpose mRNA metagene plots depicting m^6^A modifications identified through nanopore m^6^A calling and miCLIP-seq methodology, employing the Guitar R package^55^ for comparison.

### Poly A tail length estimation

The estimation of poly A tail length for each read was estimated using tailfindR v.1.3^26^. Additionally, we employed ‘nanopolish polya’ v.0.13.2^32^ to perform a comparative analysis with the tailfindR estimations. To ascertain statistically significant differences between the control and KD cells, we conducted Mann-Whitney tests (|KD – control Poly A Tail Length| > 10 (nt), FDR < 0.05). Adjusted p-values were computed using the Benjamini–Hochberg method to control the false discovery rate (FDR).

### Differential alternative splicing event analysis

We created an isoform count table using ‘FLAIR quantify’ and subsequently employed ‘FLAIR diffsplice’ through Fisher’s exact test. As a result, we identified significant alternative splicing events in two sets of control and METTL3 KD, where the p-value from Fisher’s exact test was found to be less than 0.05^56^.

### Differentially expressed isoform analysis

After aligning reads acquired through nanopore direct RNA sequencing to reference transcripts, gene expression levels were quantified using Bambu^57^. The quantification of expression was normalized as counts per million (CPM). Differentially expressed genes (DEGs) were identified through DESeq2 (Log_2_ Fold Change| ≥ 1, Adjusted p-value < 0.05)^58^.

### Correlation analysis of m^6^A frequencies and poly A tail length

We integrated m^6^A probability and poly A tail length by unique ‘read_id” identifier. The m6Anet provided ‘read_index’ information, while tailfindR offered ‘read_id’. Utilizing ‘f5c eventalign’ summary file, which included ‘read_index’ and ‘read_id’, we merged m^6^A probability and poly A tail length on the same sequencing read.

To investigate the influence of m^6^A presence on poly A tail length, we categorized read IDs into two distinct groups: ‘without m^6^A’ (m6Anet probability = 0) and ‘with m^6^A ‘ (m6Anet probability ≥ 0.6). Subsequently, transcripts with a read count ≥ 10 from each group were selected for further analysis. A Mann-Whitney test was then conducted to compare the poly A tail lengths between these two groups (|’without m^6^A ‘ – ‘with m^6^A ‘ Poly A Tail Length| ≥ 10 (nt), p.value < 0.05).

### Correlation analysis of poly A tail length and transcript expression

Poly A tail length and transcript per million normalized expression (TPM) were merged by Ensemble transcript IDs. Poly A tail lengths were derived from tailfindR, TPM was calculated using Isoquant^43^. Read counts ≥ 10 were selected for further analysis. Utilizing this merged dataset, Spearman’s correlations were computed for each Ensemble gene ID. Significant correlations (Spearman’s |R| > 0.1, FDR < 0.05) were then retrieved for Gene Ontology (GO) term pathway enrichment analysis. Genes related to ribosomal and mitochondrial genes were excluded from consideration in this analysis.

### Correlation analysis of m^6^A frequencies and transcript expression

m^6^A probability data and transcript expression (TPM) was integrated by Ensemble transcript IDs. The m^6^A probability information and transcript expression values were reported from m6Anet and Isoquant^43^, respectively. Read counts ≥ 10 were isolated for subsequent analysis. To explore alternative perspectives on two relationships, the fold change of the m^6^A modification ratio was computed, considering KD as the baseline (m6Anet probability ≥ 0.6). Additionally, the fold change of gene expression was calculated with KD as the baseline. Ensemble gene IDs exhibiting significant fold changes were depicted on quadrant diagrams (|Log_2_ Fold Change| ≥ 0.5, p-value < 0.05). These Ensemble gene IDs were categorized into four groups: ‘Hyper-methylated & Up-regulated,’ ‘Hyper-methylated & Down-regulated,’ ‘Hypo-methylated & Up-regulated,’ and ‘Hypo-methylated & Down-regulated.’

### Gene Ontology (GO) enrichment analysis

Enrichment analyses for Gene Ontology pathways were performed with clusterProfiler^59^ on genes displaying notable differences in m^6^A modification levels, poly A tail lengths, and correlation outcomes. The Biological Pathway database was utilized for this analysis. Terms exhibiting a Benjamini-Hochberg adjusted p-value of less than 0.01 were regarded as significantly enriched.

## Data accessibility

All raw dRNA-seq data were deposited to NCBI-GEO accession number SUB14548164. All analysis pipelines were made available via github.com/sunsetyerin/nanopore_dRNA_pipeline.

## Acknowledgments

The work was supported by Canadian Institutes for Health Research (CIHR) and the Japan Agency for Medical Research and Development (AMED) collaborative grant under the Canada-Japan Cooperation in Science and Technology Agreement to L.P.V. L.P.V is the Scholar of the American Society of Hematology, Scholar of the V foundation for Cancer Research and is a Tier 2 Canada Research in RNA biology and Hematological Malignancies and is supported by the Michael Smith Health Research Scholar award. L.P.V’s lab is supported by Canadian Institutes for Health Research Project Grant, Natural Sciences and Engineering Research Council of Canada Discovery Grant and the Terry Fox Research Institute New Investigator Award. We thank the staffs of the Canada’s Michael Smith Genome Sciences Centre (GSC) at BC Cancer for their technical assistance. We are grateful to our lab members for their discussion and technical assistance.

## Author Contributions

Conceptualization, L.P.V.; Methodology, Y.K., K.O.N., J.M.G., L.S., S.H.M., M.G., Y.P., S.J.M.J and L.P.V.; Formal Analysis, Y.K., K.O.N., J.M.G., K.S., S.H.M., M.G. ; Investigation, Y.K., K.O.N., L.S., S.J.M.J and L.P.V; Resources, S.J.M.J and L.P.V; Writing – Original Draft, Y.K., and L.P.V.; Writing – Review & Editing, Y.K., K.O.N., J.M.G., L.S., S.H.M., M.G., Y.P., S.J.M.J and L.P.V. Supervision, S.J.M.J and L.P.V.; Funding Acquisition, S.J.M.J and L.P.V.

## Conflict of Interest

The authors have no competing interests to declare.

## Declaration of generative AI and AI-assisted technologies

The authors declare that no generative AI or AI-assisted technologies were utilized in any aspect of this paper

## Notes

### Competing Interest Statement

The authors have declared no competing interest.

